# Beyond over- or under-sampling: autistic children’s inflexibility in sampling costly information

**DOI:** 10.1101/2024.02.04.578786

**Authors:** Haoyang Lu, Hang Zhang, Li Yi

**Affiliations:** School of Psychological and Cognitive Sciences, Peking University, Beijing, China; PKU-IDG/ McGovern Institute for Brain Research, Peking University, Beijing, China; Peking-Tsinghua Center for Life Sciences, Peking University, Beijing, China; Chinese Institute for Brain Research, Beijing, China

**Keywords:** autism spectrum disorder, information sampling, computational modeling, cognitive flexibility, Bayesian account

## Abstract

**Background:** Efficient information sampling is crucial for human inference and decision-making, even for young children. Information sampling is also closely associated with the core symptoms of autism spectrum disorder (ASD), since both the social interaction difficulties and repetitive behaviors suggest that autistic people may sample information from the environment distinctively. Previous research on information sampling in ASD focused mainly on adolescents and adults, and on whether they over- or under-sample. The specific ways in which autistic children sample information, especially when facing explicit costs and adapting to environmental changes, remain unclear.

**Methods:** We employed an adapted bead task to investigate the sampling behavior of 24 autistic and 41 neurotypical children, matched for age and IQ. In each trial of our experiment, children gathered information about an unknown target isle by drawing samples from it and then guessed the target between two isles based on their samples. In conditions where sampling was costly, children needed to weigh the benefits of information against the costs of acquiring additional samples. Through computational modeling and intricate behavioral measures, we revealed how the two groups of children differed in sampling decisions and underlying cognitive mechanisms.

**Results:** Under conditions involving costs, autistic children showed less efficient sampling than their neurotypical peers. This inefficiency was due to their increased variability in the number of samples taken across trials rather than a systematic bias. Computational models indicated that while both groups shared a similar decision process, autistic children’s sampling decisions were less influenced by dynamic changes and more driven by recent evidence, thus leading to their increased sampling variation and reduced efficiency.

**Limitations:** To refine ASD subtyping and correlate symptom severity with behavioral characteristics and computational findings, future research may need larger participant groups and more comprehensive clinical assessments.

**Conclusions:** This study reveals an inefficiency of autistic children in information sampling and tracks down this inefficiency to their increased sampling variability, primarily due to their cognitive preference for more local and static information. These findings are consistent with several influential behavioral theories of ASD and highlight the needs of a multi-level understanding of cognitive flexibility in ASD.

## Background

Imagine you’re at a newly opened gelato shop with a vast menu of flavors. While you might be tempted to inquire about each flavor to make the perfect choice, the key challenge lies not just in making the final choice, but in knowing when to stop gathering information and proceed to a decision. This scenario mirrors a critical aspect of everyday lives of everyone, including children. From basic physical laws and causality to language and social norms, children and even infants heavily rely on seeking novel information under uncertainty, especially given their limited prior experience to refer to. Previous studies have found that children and even infants are active information seekers, who further use the gathered information to guide their decisions under uncertainty [1–9]. Moreover, information often comes with a price that could be implicit (e.g., time and effort) or explicit (e.g., monetary cost). For example, children might ask questions to gain information, but they also risk experiencing embarrassment and rejection. Despite the importance of adaptive information sampling in childhood, how children modulate their sampling based on the interplay between information gain and costs has not been thoroughly examined. Researchers have only known that young children might be suboptimal than older children or adults in trading-off information gain and sampling cost [2, 5, 10]. The more intricate cognitive process underlying the adaptive information sampling in children, particularly in autistic children, is still elusive.

Autism spectrum disorder (ASD) is a neurodevelopmental disorder characterized by challenges with social interactions and restricted and repetitive behaviors [11, 12]. These core symptoms, though seemingly unrelated, are deeply intertwined with how autistic people interact with and sample information from their environment. For example, social interactions necessitate constant information sampling, such as gathering information from others’ facial expressions, tone of speech, body language, etc. to infer the mental state of others in order to keep the conversation going [13]. Repetitive behaviors could also be viewed as inefficient information sampling in reducing uncertainty due to the prolonged time spent on redundant details [14]. In a recent Bayesian account of ASD that explains a variety of perceptual, motor, and cognitive symptoms, the deviation from Bayesian optimality in information process is primary to ASD [14]. Under this Bayesian framework, information sampling is referred as “disambiguatory active inference”, whose general impairments are believed to underlie autistic symptoms. Indeed, autistic people show atypical gaze patterns in ambiguous or social scenes, sampling the visual environment in a less efficient manner [15–17].

Previous studies that directly examine information sampling in autistic people predominantly test autistic adults or adolescents. Early studies presented mixed results, with some suggesting autistic adults sample less and decide faster than neurotypical adults [18, 19], while others reporting more extensive information gathering in similar tasks [20, 21]. These discrepancies suggest that ASD may affect one’s ability to sample *optimally* dependent on particular situations, instead of oversampling versus undersampling. Using an inference task with explicit sampling costs, Lu et al. found that people with higher autistic traits have more varied and less efficient sampling behaviors, due to using a more inflexible strategy of balancing information gain and cost [22]. These studies on adults or adolescents have provided insights into autistic people’s sampling behaviors, but because information sampling may change significantly from childhood through adolescence to adulthood [2, 5, 9, 10], the information sampling behavior of autistic children is still elusive.

In the present study, we investigated the possible differences in information sampling between autistic children and preschoolers with IQ and age matched. Based on the results of previous studies, we hypothesized that autistic children would exhibit inefficient sampling when facing the complicated trade-off between information gain and cost, which might result from the varied sampling across trials. We modified the bead inference task of Lu et al. to be more child-friendly and focused on how efficiently children would adapt their strategy to various settings of evidence strength and information cost [22]. We found that autistic children had lower sampling efficiency (i.e., lower gains) under costed conditions, due to more varied sampling choices across trials rather than systematic deviation from the optimal number of samples. Computational modeling further indicates that autistic children’s lower efficiency and greater sampling variation are due to their reduced reliance on both the information from previous trials and the cost accumulated during sampling, along with a stronger influence of recently obtained information within a trial.

## Methods

### Participants

We collected data from 30 autistic children and 47 neurotypical children with ages ranging from five to eight years old. The sample size was determined by both resource limits and the typical sample size of related decision-making and learning studies in ASD [23] (see Limitations section for discussion). All the neurotypical children came from regular elementary schools. Twenty-six of the autistic children were from a special education school for autistic children in two waves and four autistic children were recruited via advertising posters in the local community. Parents of neurotypical children reported no concerns of any developmental condition. All the autistic children had received a formal diagnosis of an autism spectrum condition from pediatrics based on Diagnostic and Statistical Manual - 5 criteria [24]. As the autistic children were recruited across multiple waves and sites, fifteen autistic children in our final sample received additional assessment by the Autism Diagnostic Observation Schedule (ADOS; [25]), five by both the Childhood Autism Rating Scale (CARS; [26]) and the Autism Spectrum Quotient—Children’s Version (AQ-Child; [27]), and four only by AQ-Child (see Table 1). Nevertheless, the inclusion criteria were not solely based on meeting cut-offs of these instruments, as clinical judgment by professional clinicians or pediatrics has been shown to enhance the diagnosis stability [28]. The study abided by the Declaration of Helsinki and was approved by the Institutional Review Board at Peking University, and the study obtained oral consents from all children and written consents from their parents.

**Table 1.**
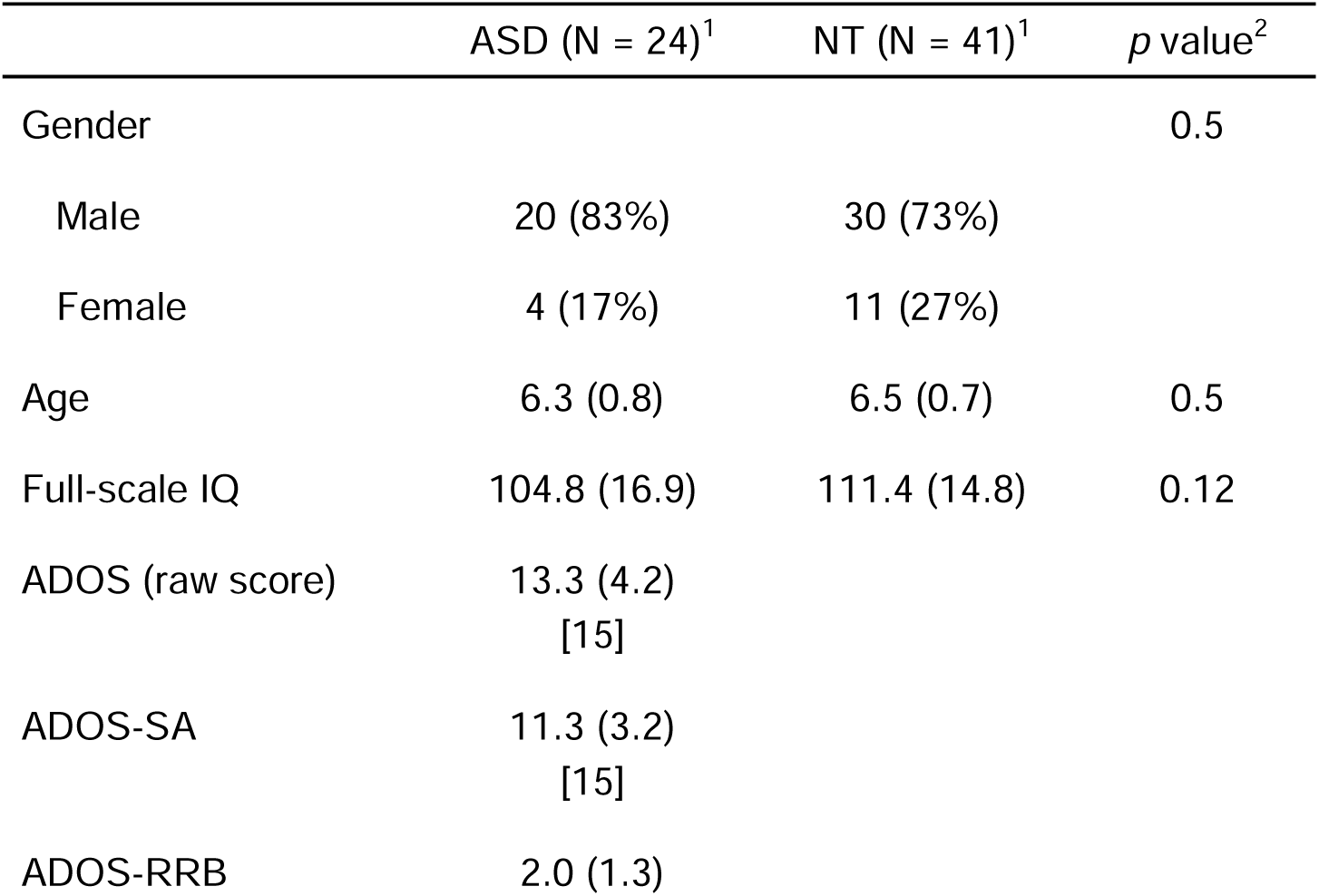
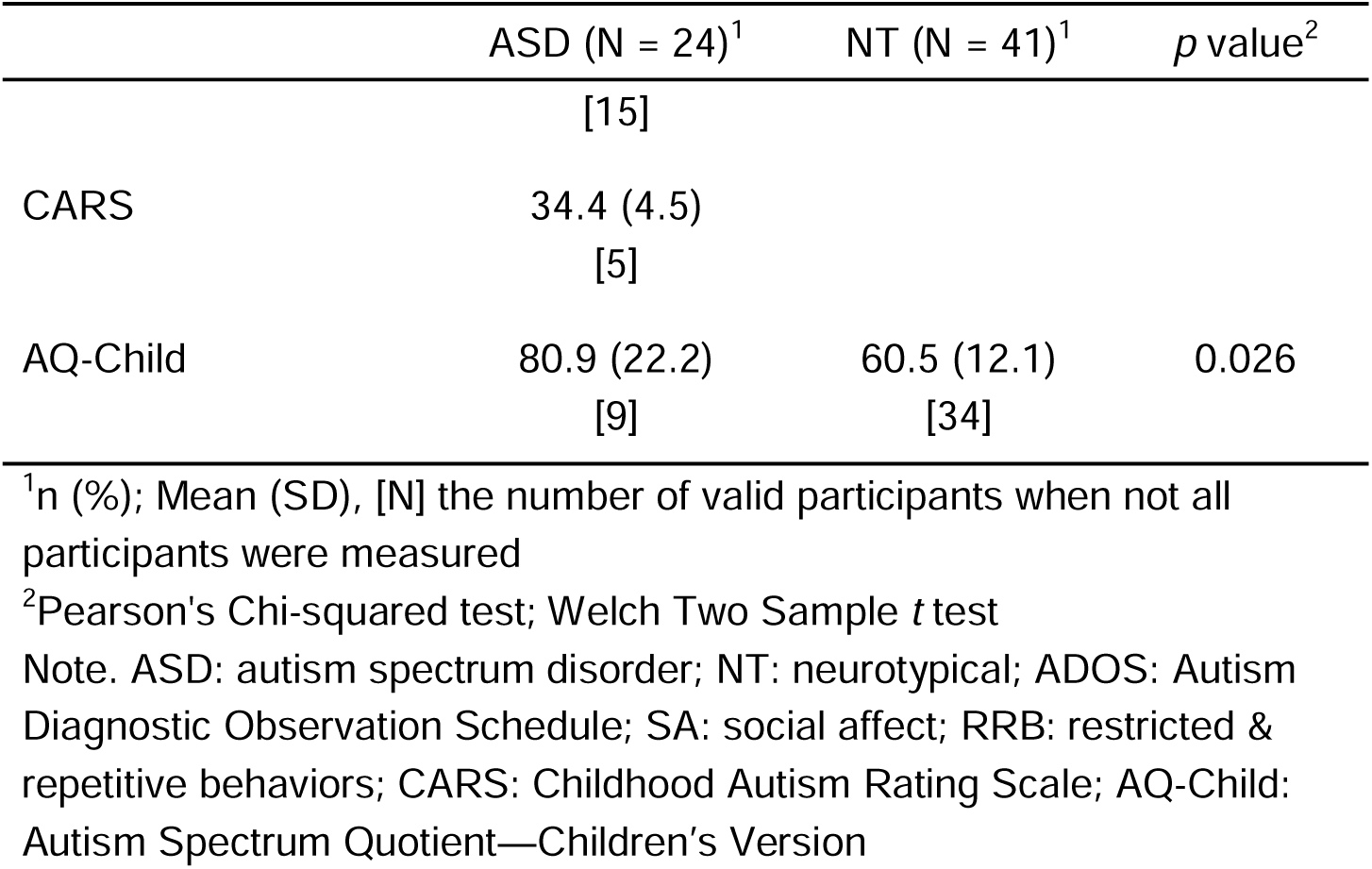
Children demographic information

Five autistic children did not understand the instructions to pass the practice trials, and thus they did not take test trials. Because of emotional distress, one autistic child quitted the experiment after completing half of the test trials, whose data were included in behavioral analyses but not in computational modeling. Six neurotypical children were not included for further analyses because of either not understanding the instruction, inattention to the task, or the experiment program crash. The final sample consists of 24 autistic children and 41 neurotypical children with two groups matched on ages and full-scale IQ (see Table 1 for detailed information). Full-scale IQ was measured by the Chinese version of the Wechsler scales [29, 30].

### Stimuli and Procedure

The task was based on the “bead task”, a classic instrumental information sampling task [31, 32], and adapted as an adventure game to be of interest to the children (Figure 1a). On each trial, children saw two isles on both sides of the monitor screen. Two isles were habited by ten animals with the opposite ratio of dogs to cats, either 60% to 40% vs. 40% to 60% in a block of trials or 80% to 20% vs. 20% to 80%. Children were told that the goal on each trial was to find out on which isle they were on, based on the animals they had met. They were free to encounter 20 animals at most by pressing the button on an Xbox controller, and when they felt confident enough to make the decision, they could press the corresponding buttons to choose the left or right isles shown on the screen. They would receive 100 credits for correct responses and zero for incorrect responses. In some conditions of the experiment, it would cost children 0, 1, or 4 credits for each animal, in which case the actual gain for a correct response was 100 credits deducted by the cumulative cost incurred for finding animals; an incorrect response would still yield zero credit. At the end of the experiment, the experimenter would give children several of their favorite stickers in proportion to their accumulated credits across trials. Children were encouraged to sample (i.e., choose to encounter animals) wisely to maximize their game credits and final rewards, which would require proper trading-off between information gain and cost during the task (Figure 1b).

**Figure 1.**
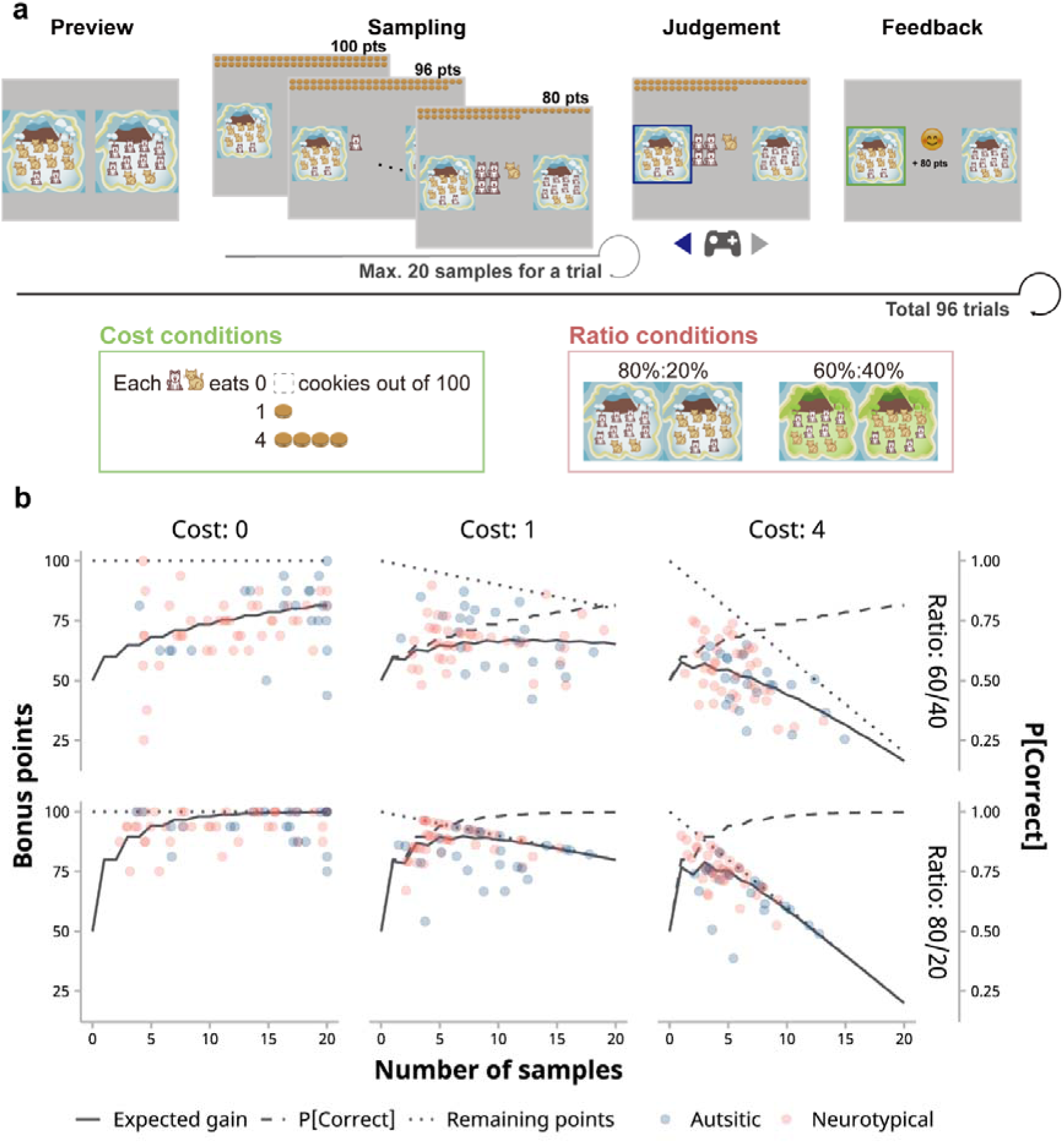
Information sampling “bead” task. (a) The child-friendly “bead task” substituted beads with cat and dog images to enhance engagement. Each block began with a preview of the cost and ratio conditions for subsequent trials (“Preview”). During a trial, children might sample up to 20 animals from a predetermined isle by pressing a button on the gamepad (“Sampling”). When children wanted to make a judgement, they could choose by pressing the left or right button on the gamepad (“Judgement”). The task was structured into six blocks, each a combination of three cost conditions and two ratio conditions. (b) Children typically faced a trade-off between information gain and cost: the expected probability of a correct judgement (dashed lines) increases, while the remaining bonus points (dotted lines) decrease as a function of the number of samples. Consequently, the expected gain, a product of these factors, initially increases but then decreases (as shown by solid lines, especially noticeable in the “Cost: 1” and “Cost: 4” panels). Each point denotes the actual points won by each child averaged across trials.

The experiment consisted of three blocks corresponding to three cost conditions (Figure 1a; zero-cost: 0, low-cost: 1, or high-cost: 4 credits). The order of three blocks was counterbalanced across children. Two mini-blocks of different dog-to-cat ratios (low-evidence: 60%:40% and high-evidence: 80%:20%, respectively) were nested under each block in a random order. For most of the children, each mini block had 16 trials, so the experiment had 96 trials in total. However, because some autistic children were more prone to feel upset during the long task, six autistic children participated in a shorter version of the experiment with 8 trials in each mini block (i.e., 48 trials in total).

To ensure children had requisite understanding of the task, the experimenter asked a series of questions about the ratios, costs, and rewards after the introduction (see Additional File 1, S1 Text for the task instruction and comprehension questions). Only after correctly answering all the questions could children enter the practice task. The practice task was a mock test having only 12 trials but the same structure as the main task. To pass the practice, children had to be “theoretically correct” in more than 75% of the trials: being “theoretically correct” was to choose the isle where the majority of animals was matched with the majority of encountered animals, regardless of the actual outcome. For example, if children have met more dogs than cats, they should choose the dog isle because of higher likelihood of being correct. If children failed to pass the practice, the experimenter would repeat the instruction and the practice task for them. However, if children failed the practice again, they were not eligible to start the main task.

The stimuli were presented on a 24-inch monitor during the experiment via Psychtoolbox in MATLAB 2016b. Children sat about 60 cm in front of the monitor, holding an Xbox controller. We chose dog and cat icons that were similar in style and the same in size, average brightness, and contrast to mitigate the choice bias caused by familiarity or visual salience. To ease the difficulty of memorizing all related information for children, the ratio and cost information would be displayed before the beginning of the next mini-block, and the experimenter would repeat the information to them as well. Additionally, the remaining credits and encountered animals would always be shown during a trial so that children could know their accumulated costs and evidence at any time.

### Statistical analyses

All statistical analyses were conducted in R 4.3.2 [33]. We used linear mixed models for model-free analyses to account for the imbalanced data resulting from the unequal numbers of children and trials. Linear mixed models were conducted on the trial level, which utilized more information of the data than condition average-based repeated measures ANOVA. Linear mixed models were estimated using “afex” package [34], and *F* statistics, degrees of freedom of residuals (denominators), and *p*-values of the models were approximated by Kenward-Roger method [35, 36]. Specifications of random effects followed parsimonious modeling [37]. For significant fixed effects, “emmeans” package was used to test follow-up simple effects and post hoc contrasts [38]. Statistical multiplicity of the post hoc contrasts was controlled by a single-step adjustment that used multivariate *t* distributions to estimate the critical value for the contrasts [39].

Specifications of (generalized) linear mixed models for behavioral data are set as followed (using Wilkinson notation, [40]; * represents main effects of predictors and interaction effects between all predictors plus all lower-order terms):

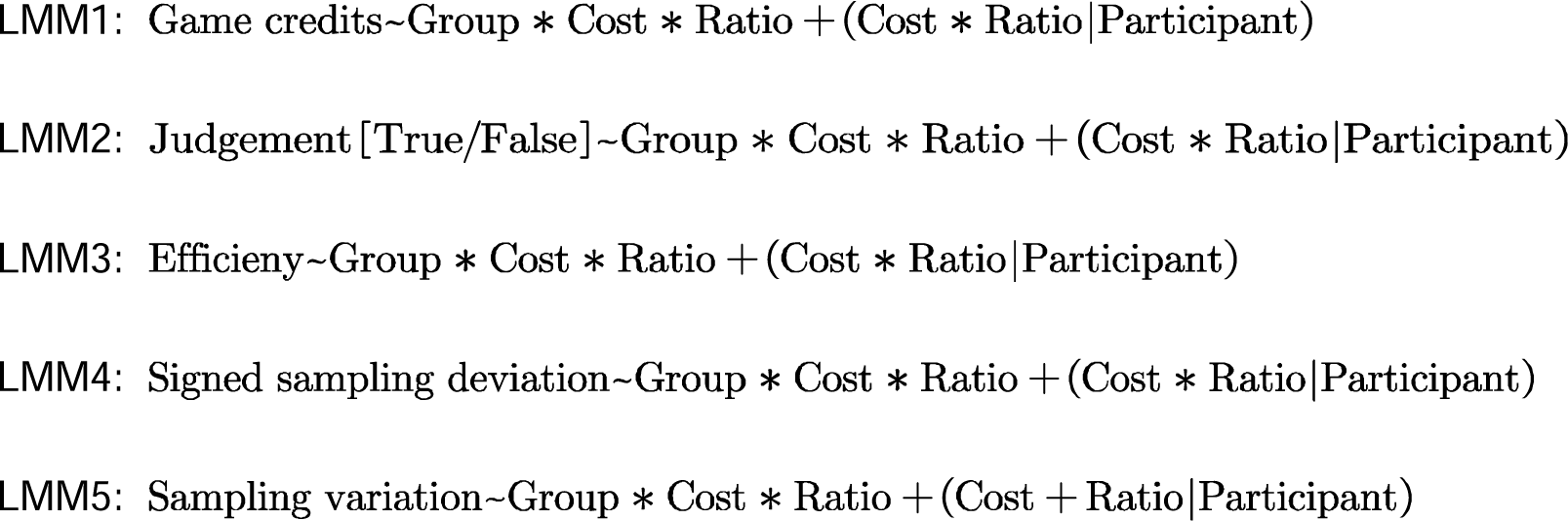

Efficiency in LMM3 is the same as in Lu et al., which is the expected gain for children’s animal sample sizes divided by the maximum expected gain given the sampling cost and ratio condition [22]. Suppose the animal sample size is *n*, the maximal reward is 100 credits, the unit sample cost is *c*, and the percentage of predominated animals in the preselected isle is *q*, then the expected gain is 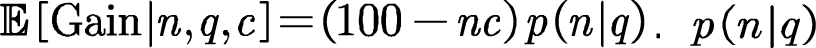 is the expected probability of correct judgment, defined as follows:

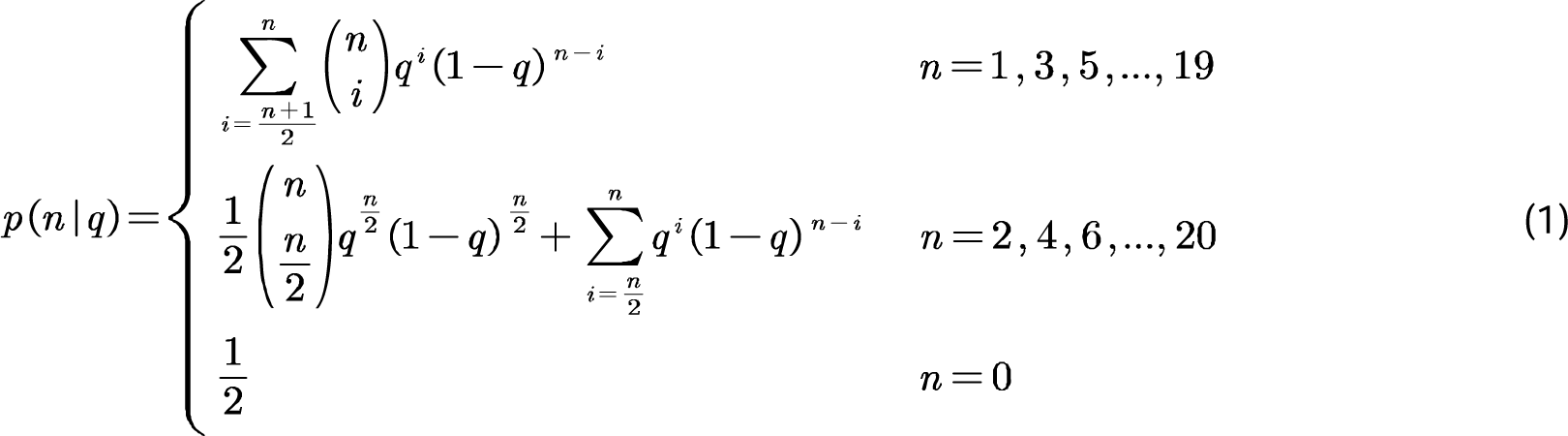

Thus, the optimal sample size is the value of *n* that maximizes 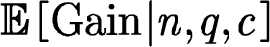 To investigate the sampling strategy that might result in inefficient sampling, we further calculate signed sampling deviation (LMM4), which is the signed sample size difference from the optimal sample size, and sampling variation (LMM5), which captures the inter-trial standard deviation of sample sizes in a condition.

### Behavioral modeling

To account for the difference in the cognitive mechanism behind the sampling decision of children, we adopted six computational models described in Lu et al. [22]. These six models could be divided into two one-stage models and four two-stage models (see Figure 3a). In short, one-stage models assumed the decision of stop sampling was determined by the linear combination of decision variables (such as cost-related and evidence-related variables) at once, whereas for two-stage models, the decision of stop sampling was first determined by a part of decision variables and probabilistically given a second thought to consider other variables. In one-stage models, the probability of stopping sampling on the *i*-th trial after having drawn *j* samples is as a function of the linear combination of *K* decision variables (DVs) via a logistic function:

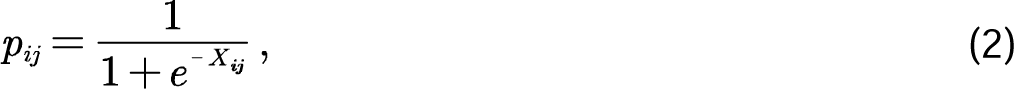

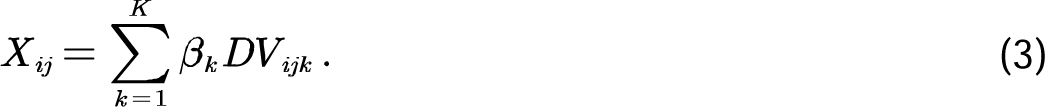

Four one-stage models differed in the combination of decision variables:

1. Cost-only model (denoted Cost only): cost-related variables, including unit cost per animal sample (categorical: 0, 1, 4; 0 was set as the reference level), number of animals sampled, and total sampling cost (product of the former two terms),
2. Cost and evidence with decay model (denoted Cost + Evidence): both cost-related and decayed evidence-related variables. On top of Cost only model, Cost + Evidence further includes unit log evidence per animal sample (i.e., 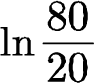 or 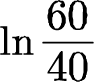), absolute value of cumulative information (cumulative information refers to the difference between the numbers of “cat” and “dog” samples), total log evidence (product of the former two terms), and the correctness and the number of animal samples in the last trial. In models with decayed evidence, cumulative information (CI) is modulated by a decay parameter α:

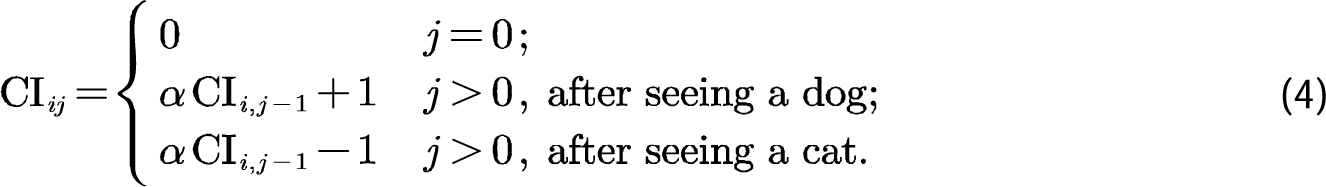 A value of the decay parameter near 0 indicates a tendency to consider only the most recently obtained information, whereas a value closer to 1 would mean considering all evidence with nearly equal weight. In two-stage models, sampling choices may involve two decision stages, with the probability of reaching the decision of stopping sampling in each stage being

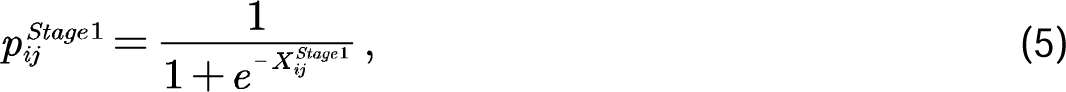

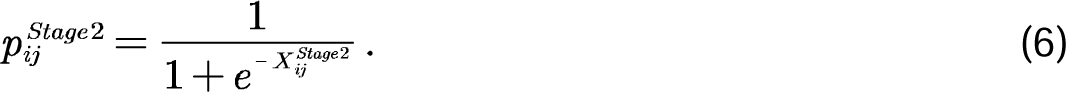 The overall probability of stopping sampling can be written as

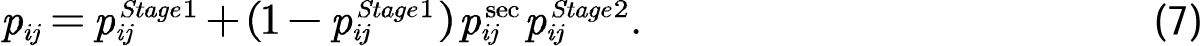 Four second-stage models differed in both the combination of decision variables and the second-thought probability, including:
3. 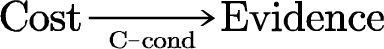: the first stage and the second-thought probability were both determined by cost-related variables, whereas the second stage was determined by evidence-related variables,
4. 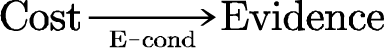: the first stage was determined by cost-related variables, but the second-thought probability was controlled by evidence conditions,
5. 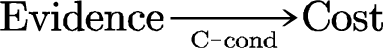: the first stage was determined by evidence-related variables, but the second-thought probability was controlled by cost conditions,
6. 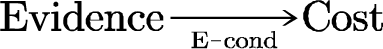: the first stage and the second-thought probability were determined by evidence-related variables.

When the second-thought probabilities were conditional on cost conditions, *p_ij_*^sec^ = *p*_zero_^cost^, *p*^low cost^, and *p*^high cost^ respectively for the zero-, low-, and high-cost conditions; when conditional on evidence conditions, *p_ij_*^sec^ and *p*^high evidence^ respectively for the low- and high-evidence conditions. In general, model parameters to be estimated included β_k_ that represents the effects of decision variables, decay parameter α_Decay_, and second-thought probabilities *p_ij_*^sec^ if two-stage models (see Addition File 1, S1 Table).

### Model fitting and comparison

We fitted models separately to the behavioral data of neurotypical and autistic children, using hierarchical Bayesian estimation with Hamiltonian Monte Carlo implemented in Stan and cmdstanr package in R [41, 42]. In contrast to Lu et al. using maximum likelihood estimation, the hierarchical Bayesian estimation allows incorporation of prior information and sharing information across participants, which is especially beneficial for relatively small sample sizes and provide more informative and robust results.

Each of four separate Markov chains with randomized initial values took an adaptive warm-up with 5000 samples from the posterior to prevent the dependence on the initial values, and then each chain took another 10000 samples. This resulted in 40000 samples from the posterior for each parameter. For complex hierarchical models, in accordance with the standard practice in Stan, parameters were sampled using non-centered parameterization; that is, parameters were independently sampled from a standard normal distribution before transformed to an appropriate range. Convergence between chains was confirmed based on Gelman-Rubin *R̂* of all parameters being less than 1.01 and no systematic divergent transitions. All individual-level parameters were assumed to sampled from the corresponding group-level normal distributions, whose mean and standard deviation were estimated from the data (parameter specifications see Additional File 1, S1 Text).

Model comparison was performed based on the pointwise out-of-sample prediction accuracy of the model, which was evaluated by the Bayesian leave-one-out cross validation estimate of expected log predictive density (LOO ELPD) and implemented in “loo” package in R [43, 44]. Moreover, models could be compared in terms of the model weights from pseudo-Bayesian model averaging with Bayesian bootstrapping (Pseudo-BMA+; [45]). The model weights were calculated taking into account the penalty for model complexity. Models with higher ELPD or Pseudo-BMA+ model wights suggest better performance while having a parsimonious model specification. All models were compared within each group of children (i.e., in autistic and neurotypical children separately).

## Results

### Autistic children differed from neurotypical children in sampling strategy rather than correct judgement

Overall, autistic children took fewer samples than neurotypical children overall (Figure 2a; *M*_ASD_ = 10.3, *M*_NT_ = 7.6, *F*(1, 63.17) = 11.03, *p* < .001), but their gain was not significantly fewer than neurotypical (NT) children (LMM1; *M*_ASD_ = 72.4, *M*_NT_ = 74.9, *F*(1, 63.32) = 3.90, *p* = .053). However, when we examined different conditions separately, we found that autistic children had won significantly fewer credits in the high-cost condition (*M*_ASD_ = 57.4, *M*_NT_ = 64.7, *t*(66.6) = –3.14, *p* = .007) and in the high-evidence condition (*M*_ASD_ = 80.3, *M*_NT_ = 85.7, *t*(182.7) = –4.00, *p* < .001). An analysis of the proportion of correct judgment allows us to exclude the possibility that autistic children made more incorrect judgement. There was no significant difference between the two groups in the proportion of correct judgement (LMM2; _χ_ (1) = 0.33, *p* = .57). Though a significant interaction effect was found between the group and the evidence conditions (LMM2; ^2^(1) = 5.08, *p* = .024), post hoc comparisons did not support significant group differences under any of the two evidence conditions (low-evidence condition *OR* = 1.16, 95% CI: [0.92, 1.47], *p* = .28; high-evidence condition *OR* = 0.73, 95% CI: [0.46, 1.18], *p* = .26). When comparing two groups in terms of the gained credits in those correct trials, we found that autistic children won fewer credits particularly in those costly trials (interaction between groups and cost conditions: *F*(2, 67.30) = 4.07, *p* = .021; low-cost condition: *t*(63.4) = -2.58, *p* = .033; high-cost condition: *t*(63.2) = -2.68, *p* = .026). These results suggest that the main behavioral difference was pertinent to their sampling strategy rather than judgement.

**Figure 2.**
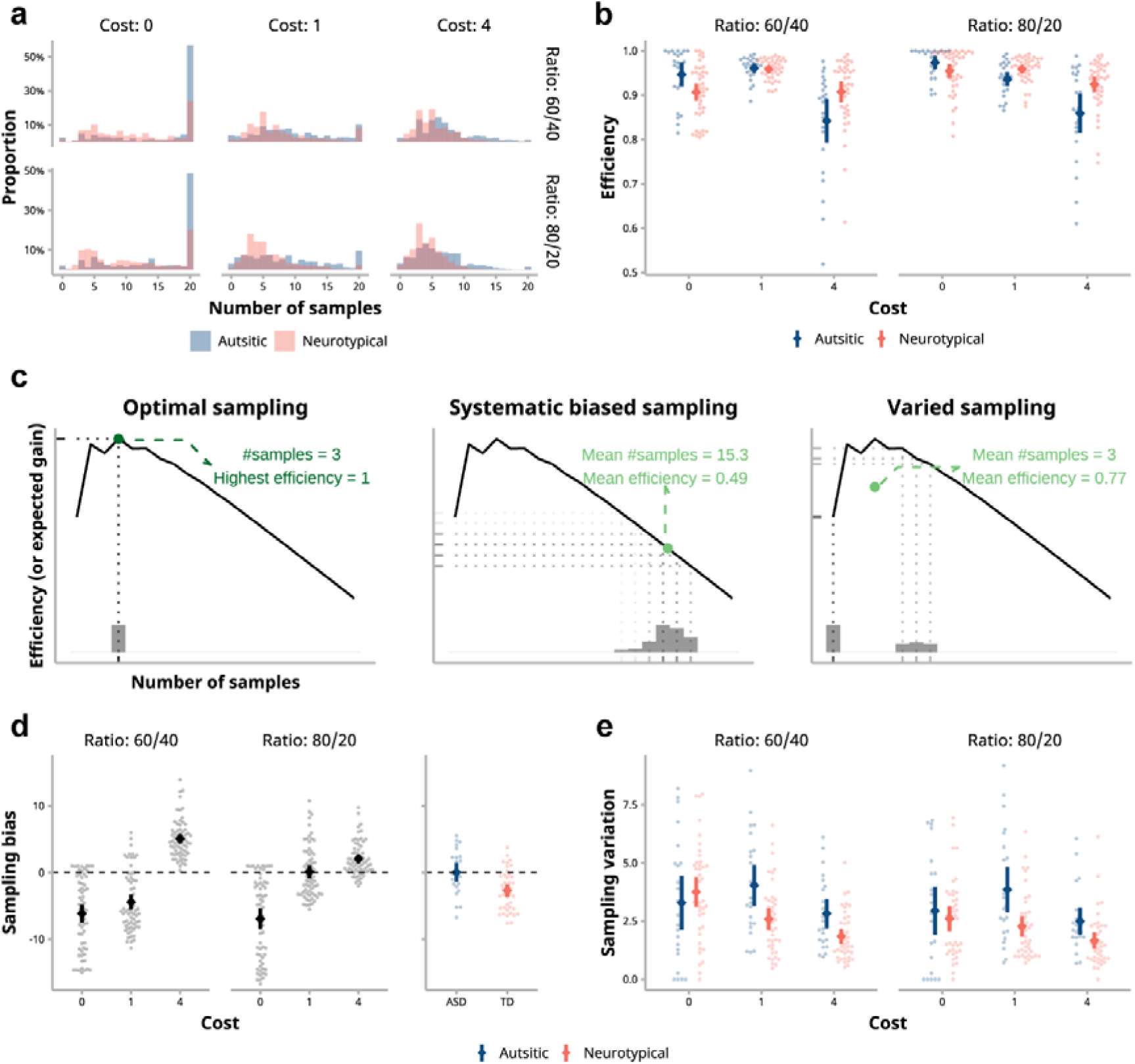
Sampling performance of autistic and neurotypical children. (a) Distributions of the average number of samples of each child. Children took fewer samples as the sampling cost or sample ratio increased. Autistic children in general sampled more than neurotypical (NT) children. Histograms of two groups are overlaid with one another. (b) Sampling efficiency is the ratio of expected gain from a given number of samples to the maximum expected gain from the optimal sample number. The sampling efficiency of autistic children was significantly lower than that of NT children, and the difference was even larger in the high-cost condition. (c) When children could stick to the (near-)optimal sample (three samples, in the case of the figure), they would achieve the highest efficiency (left panel). However, both systematic bias and large variation in sampling would lead to poorer efficiency (middle and right panels). Notice that even when the mean number of samples equates to the optimal, that individuals sometimes take too few samples and at other times too many would nevertheless reduce the efficiency (i.e., varied sampling, right panel). Solid lines are the expected gain as the function of the number of samples, illustrating the case under the high-cost and high-evidence condition. Histograms are example distributions of samples. (d) The sampling bias was modulated by sampling cost and evidence. Autistic children did not show systematic bias, whereas NT children showed undersampling behavior in general. The difference between the two groups was significant. (e) The sampling variation of autistic children was significantly greater than that of NT children and was also moderated by the amount of costs. Under costly conditions, the sampling variation of the autistic children was greater. In (b), (d), and (e), large points and error bars are means and 95% confidence intervals, and each small point represent the mean of each child.

### Sampling behaviors: Autistic children had significantly lower efficiency than neurotypical children

An adaptive sampling strategy was used to reach the optimal decision by simultaneously considering and balancing the sampling costs and the information gained from sampling (see Figure 1b). We adopted the sampling decision efficiency to measure the optimality of children’s information sampling behavior (LMM3). Efficiency was calculated as the expected gain for the number of samples taken by participants divided by the maximum expected gain. First, we looked at the effects of experiment conditions and found a significant interaction between sampling cost and evidence (Figure 2b; *F*(2, 65.02) = 30.77, *p* < .001). Both autistic and NT children had higher sampling efficiency in high-evidence conditions than low-evidence conditions when sampling was costless (*t*(63.7) = 6.63, *p* < .001), whereas children had better performance in the low-evidence conditions than the high-evidence conditions in those low-cost trials (*t*(64.6) = -3.07, *p* = .010).

Overall, the sampling efficiency of autistic children was marginally significantly lower than that of NT children (*M*_ASD_ = 92.0%, *M*_NT_ = 93.5%; *F*(1, 63.26) = 3.96, *p* = .051). The significant interaction between group and cost suggests that the group difference increased with increasing cost (Figure 2b; *F*(2, 63.35) = 6.65, *p* = .002): there was no significant difference between the two groups in zero- and low-cost trials (*t*(63.3) = 2.26, *p* = .75, and *t*(62.9) = -1.55, *p* = 0.31 respectively), whereas in high-cost trials, the sampling efficiency of autistic children was lower than that of NT children (*t*(63.0) = -3.17, *p* = .007). We also found a significant interaction between group and evidence level (*F*(1, 63.50) = 4.28, *p* = .043) that autistic children had a significantly lower efficiency than NT children in high-evidence conditions (high-evidence: *t*(63.2) = -2.63, *p* = .019; low-evidence: *t*(63.5) = -0.97, *p* = .50).

### Sampling behaviors: Autistic children’s lower sampling efficiency came from their higher sampling variation

In the above analysis of sampling efficiency, we found that the differences between the two groups were modulated by sampling cost: the higher the cost, the worse autistic children performed relative to NT children (Figure 2b). What erred in autistic children’s sampling strategy? Greater systematic biases (i.e., oversampling or undersampling) was one possibility, but not the only one. According to our simulations, higher sampling variation may also lead to lower efficiency (see Figure 2c). Or, both bias and variation may play their roles. To elucidate these different possibilities, we decomposed children’s sampling behaviors into biases from optimality (Figure 2d) and variation (Figure 2e).

In general, children exhibited both oversampling and undersampling behavior under different conditions (LMM4; Cost x Ratio interaction: *F*(2, 62.89) = 250.12, *p* < .001): for low-evidence conditions, both groups undersampled in zero- and low-cost trials while oversampled in high-cost trials. For high-evidence conditions, children undersampled in zero-cost trials, while they oversampled in high-cost trials; in the low-cost trials, the number of samples taken by both groups was close to optimal (Figure 2d). Autistic children did not exhibit a systematic sampling bias and their signed sampling bias from the optimal was not significantly different from zero (*M* = -0.004, *t*(63.6) = -0.006, *p* = 0.99). In contrast, the NT group exhibited significant undersampling, with a sampling bias significantly less than zero (*M* = -2.72, *t*(62.4) = -5.49, *p* < .001). The difference of sampling bias between the two groups was significant (*F*(1, 63.12) = 11.03, *p* = .001). No significant interaction effects involved groups.

The analysis of sampling variation revealed a significant main effect of group, with autistic children showing greater variation in the number of samples taken across trials (Figure 2e; LMM5: *F*(1, 63.70) = 6.87, *p* = .011). There was also a significant interaction between group and sampling cost (*F*(2, 63.38) = 7.73, *p* = .007), with the variation of the autistic group being particularly high in low- and high-cost conditions compared to the NT group (*t*(64.2) = 3.72, *p* = .001 and *t*(75.4) = 2.96, *p* = .012, respectively). No other interaction effects including group predictor were significant. Considering both sampling bias and sampling variation, it appears that the high variation in sampling across trials in costly conditions may be a major cause of the lower sampling performance observed in autistic children in those conditions.

### Computational modeling: Autistic children were less influenced by dynamic and global information when making sampling decisions

To understand the cognitive mechanisms behind sampling decisions in autistic and neurotypical children, we applied models from Lu et al. [22], which were four first-stage models and four second-stage models (Figure 3a). The first-stage models described what information children integrated to achieve the sampling decision. The second-stage models further distinguished how they made the decision in two steps, with different information considered in each step. The information that the children were assumed to integrate could be categorized into cost-related (e.g., cumulative costs during sampling) and evidence-related decision variables (e.g., cumulative log evidence; Figure 3b). Using hierarchical Bayesian models, posterior probability distributions for each parameter for each child at the individual level and the entire group at the population level were obtained simultaneously. Model comparisons were performed based on leave-one-out expected log predictive density (LOO ELPD) and pseudo-Bayesian model averaging weights using resampling method (Pseudo-BMA+; see Methods for details).

**Figure 3.**
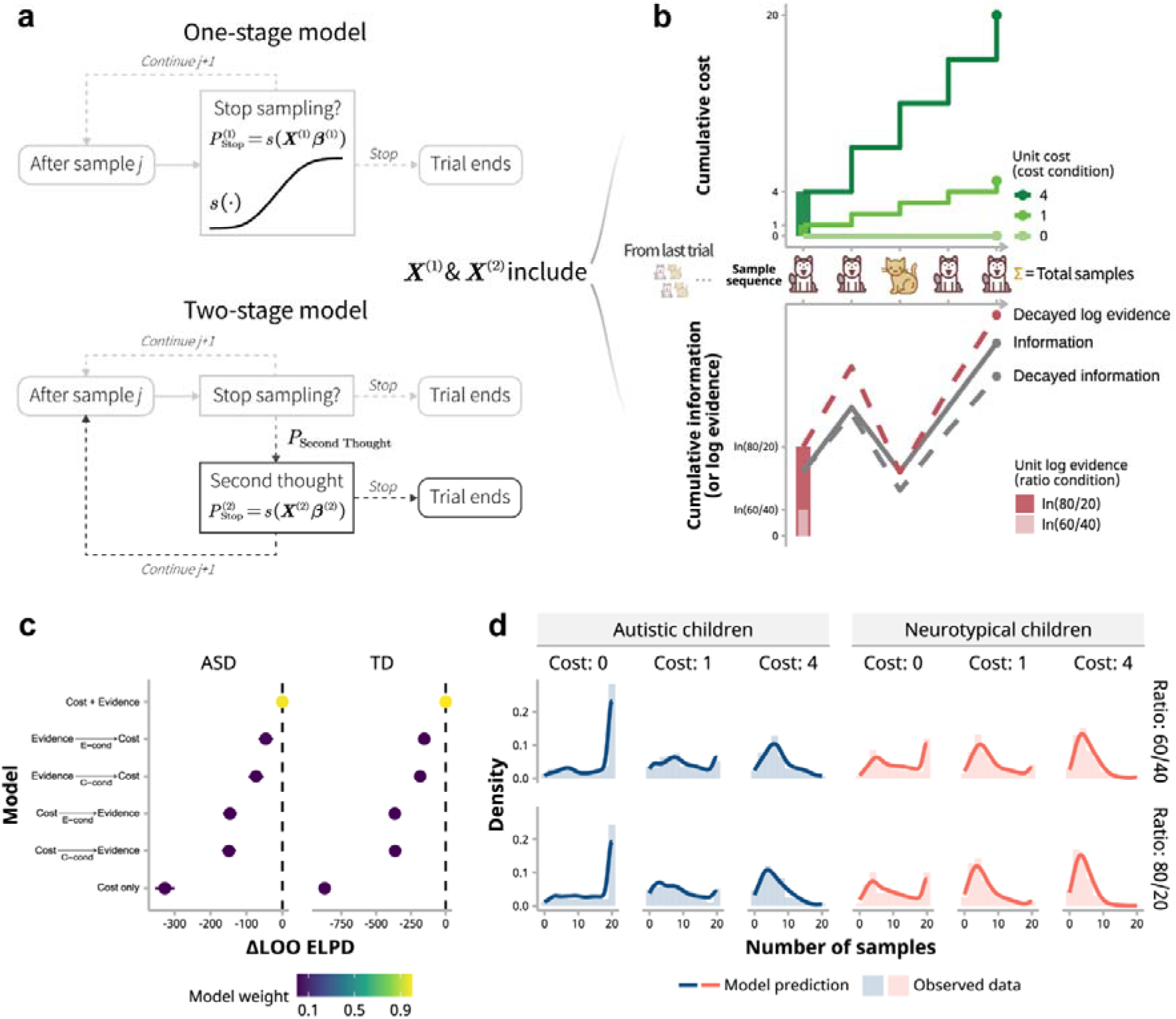
Computational modeling of sampling choices in autistic and neurotypical children. (a) Diagram of one-stage and two-stage models. The one-stage model consists of the steps shown on the top: Each time a child opts to stop sampling, the probability of doing so is determined by a sigmoid function of a linear combination of various decision variables. Additionally, the two-stage model includes a potential second stage (on the bottom), where children might re-evaluate their decision to stop sampling via the second-thought probability, and the second decision is influenced by a different set of decision variables. (b) Decision variables are categorized into cost-related and evidence-related groups. Cost-related variables encompass unit cost, total samples, and cumulative cost (primarily in the upper sub-panel), whereas evidence-related variables consist of unit log evidence, total decayed information, cumulative decayed log evidence, along with the sample number and correctness of the last trial (primarily in the lower sub-panel). (c) LOO ELPD (leave-one-out expected log predictive density) signifies the model evidence, with higher values indicating a better fit to the data. ΔLOO ELPD represents the difference in LOO ELPD relative to the best model (i.e., Cost + Evidence), where a value nearer to zero suggests a closer alignment with the best model. Error bars denote the standard errors of the LOO ELPD differences. Point colors reflect model weights assigned using the Pseudo-BMA+ method; heavier weights suggest a better model fit. (d) Comparison between the best model’s predictions and the actual data shown as the distribution of sample numbers under various conditions. The model accurately predicts the sampling behavior of both groups, with solid lines depicting observed data and histograms indicating model predictions.

In both groups of children, LOO ELPD and Pseudo-BMA+ model weights indicated that Model Cost + Evidence had the highest model evidence and was the best fit for the sampling behavior data based on the number of samples (Figure 3c). In the autistic group, LOO ELPD for Model Cost + Evidence was - 4134.68 and Pseudo-BMA+ weight for Model 4 was 0.987; in the NT group, the LOO ELPD was -7777.16 and the Pseudo-BMA+ weight was greater than 0.999. In addition, based on the estimated model and the complete stimulus sequence during the experiment (including unpresented “cat” and “dog” stimuli that would have been shown if children had not stopped sampling), we can predict the sampling choices of children. The posterior predictive checking confirmed that Model Cost + Evidence was robust to predict decision choices well for both groups of children (Figure 3d). As the best model Cost + Evidence suggested, both the autistic and NT children tended to combine both the cost-related and the evidence-related decision variables all together to determine the sampling choice.

To explore the computational mechanisms that lead to differences in behavioral outcomes between autistic and NT children, the population-level parameters of the winning Model Cost + Evidence were compared between the two groups (this method of between-group comparison at the population level is similar to that used in [46]; Figure 4).

**Figure 4.**
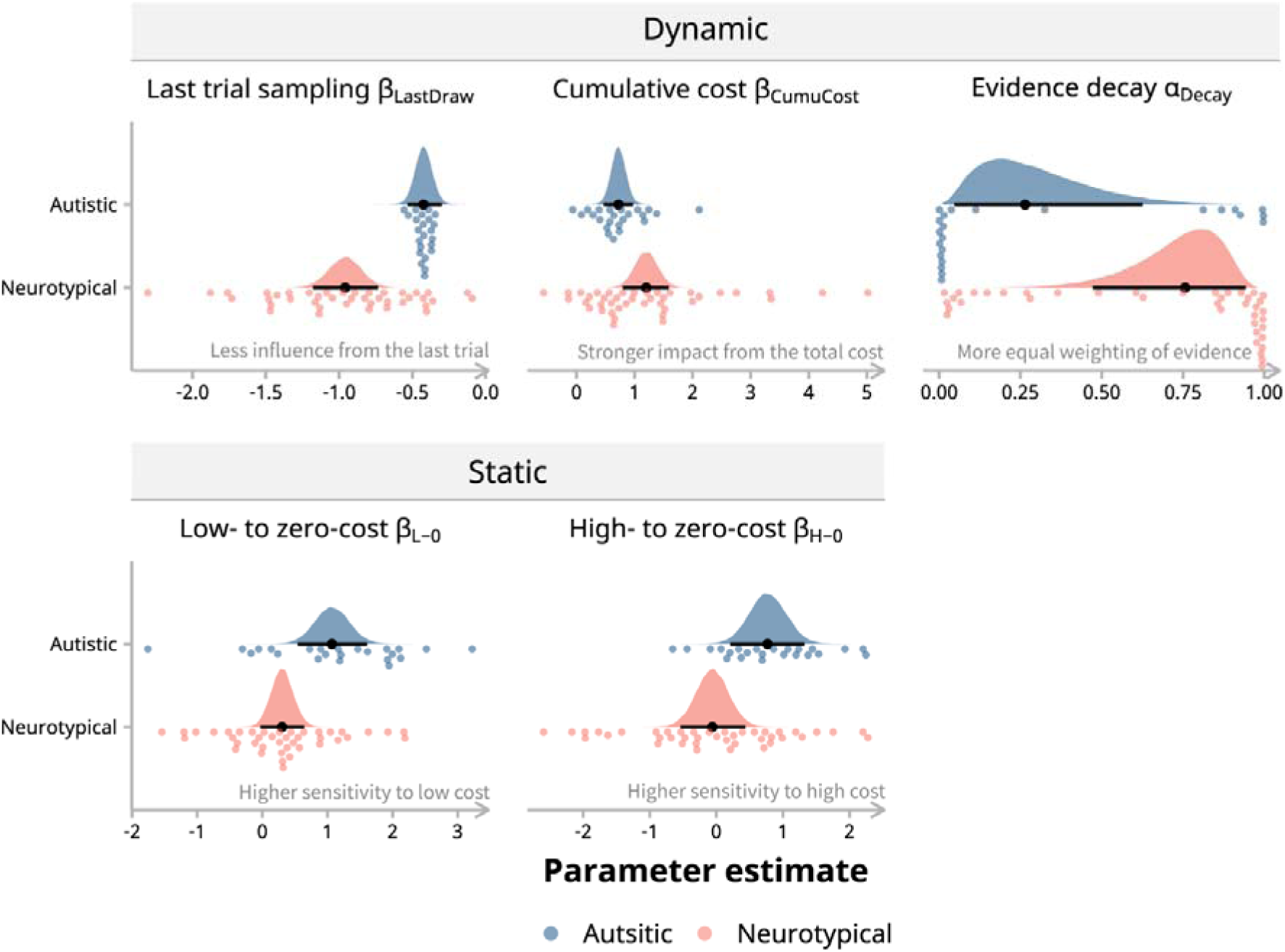
Parameter estimates from the best model in autistic and neurotypical children. Two groups of children showed significant difference in five parameters from the best model. These parameters fall into two categories that reflect dynamic and static influences of decision variables on when stop sampling. The top row (“Dynamic”) includes parameters that gauge behavioral adaptability: β_LastDraw_ represents the influence from the number of samples of the last trial, β_CumuCost_ measures the impact of the total cost accrued during the current trial, and α_Decay_ determines how information gathered during a trial is weighted. These parameters depict a flexible, adaptive process as they change with the actions taken across and within trials. The bottom row (“Static”) captures cost sensitivity: β_L-0_ and β_H-0_ indicate the general inclination to stop sampling in costly conditions compared to a zero-cost condition. These two parameters are considered “static” because they remain consistent across trials within each cost context. Overall, compared with neurotypical children, the autistic group demonstrated a reduced adaptation to dynamic shifts within the task but heightened sensitivity to the static factor of “cost as a condition”. Data points denote individual parameter estimates, shaded areas represent group-level parameter distributions, and black markers with lines illustrate medians and 95% highest density intervals (HDIs) for these distributions.

Results showed that the two groups of children differed in the coefficients of the following decision variables: cost condition differences (β_H-0_ for high-cost relative to zero-cost condition, and β_L-0_ for low- to zero-cost difference), cumulative sampling cost (β_CumuCost_), the number of samples drawn in the last trial (β_LastDraw_), as well as in the *decay* term of accumulated evidence (α_Decay_). The rest of the model parameters did not show noticeable group difference (for detailed summary statistics of two groups and the difference see Additional File 1, S2 Table).

In addition to the cost- and evidence-related taxonomy, these model parameters can also be classified as “static” and “dynamic”, by the nature of their coupled decision variables. This classification offers another perspective to understand the cognitive process differences between groups. Static parameters include β_L-0_ and β_H-0_ since they parallel the fixed differences between experimental conditions, which do not vary along with the sampling behavior. In contrast, dynamic parameters, such as β_CumuCost_, β_LastDraw_, and α_Decay_, influence sampling decisions dynamically with each sample and trial. For the static parameters, the median group difference *Mdn*_ASD_ − *Mdn*_NT_ for β_H-0_ was 0.83, 95% highest density interval (HDI) [0.11, 1.59], and for β_L-0_ it was 0.77, 95% HDI [0.16, 1.41]. For the dynamic parameters, the median differences were β_LastDraw_ at 0.54, 95% HDI [0.28, 0.78], β_CumuCost_ at -0.48, 95% HDI [-0.95, -0.01], and α_Decay_ at -0.46, 95% HDI [-0.81, -0.04]. Then, we further examine how these static and dynamic parameters affect the sampling behavior of autistic and NT children.

### Reduced adaption to dynamic information led to increased sampling variation in autistic children

First, we focused on the effects of the static parameters, β_H-0_ and β_L-0_, on sampling decisions. The parameters determined the average tendency to stop sampling under different cost conditions, as evidenced by the differences in average sample numbers between these conditions. For the autistic group, both β_H-0_ and β_L-0_ were higher compared to the NT group, indicating a heightened sensitivity to cost of autistic children. Specifically, in costly conditions, autistic children reduced their sampling more significantly (the number of samples: *M*_H_ − *M*_0_ = -8.33, *t*(63.4) = -8.44, *p* < .001; *M*_L_ − *M*_0_ = -5.64, *t*(62.8) = -6.00, *p* < .001; see Figure 2a), in contrast to NT children who showed a less pronounced decrease in sampling under the same conditions (*M*_H_ − *M*_0_ = -6.04, *t*(62.1) = -8.15, *p* < .001; *M*_L_ − *M*_0_ = -4.02, *t*(61.8) = -5.72, *p* < .001). However, these static parameters alone could not explain why the sampling variation differed between the groups since they predominantly govern the general sampling tendency within a given condition rather than influencing individual trial variations.

Therefore, we turned to the dynamic parameters, β_LastDraw_, β_CumuCost_, and α_Decay._ These parameters reflected a more dynamic and flexible cognitive process, because they would affect sampling decisions within each trial based on the previous trial or the sample history within a trial. To better illustrate their influence, we conducted simulations based on the best-fitting model and by varying interested parameters one by one within the range estimated from the children’s date while keeping other parameters fixed (Figure 5a-c). For β_LastDraw_, autistic children showed values that were larger and closer to zero compared to neurotypical children, suggesting they were less influenced by the sample numbers from the last trial in their current sampling decisions. From the equation of calculating the stopping probability (Equations 1 & 2), a more negative β_LastDraw_ value indicates a decreased likelihood of stopping sampling as the previous trial’s sample number increases, potentially leading to more samples in the current trial. Conversely, a β_LastDraw_ value closer to zero means the previous trial’s sample count has little to no impact on the current decision, allowing for greater variation in sample numbers from trial to trial, which is consistent with the simulation (Figure 5a). For β_CumuCost_, a higher value indicates that children are more likely to stop sampling as the cumulative cost increases. This means that as children encounter higher costs, they become more cautious, reducing the number of samples they take (Figure 5b). In the case of α_Decay_, a value approaching one suggests that children consider all pieces of evidence more uniformly when deciding to stop sampling. A balanced α_Decay_ (i.e., closer to one) means the decision to sample is less swayed by the recency or initial pieces of evidence, potentially leading to a more consistent number of samples across trials. This is more evident in zero-cost conditions and high-evidence conditions where sampling decisions should not be limited by the cost and the evidence is easier to accumulate in one direction (Figure 5c).

**Figure 5.**
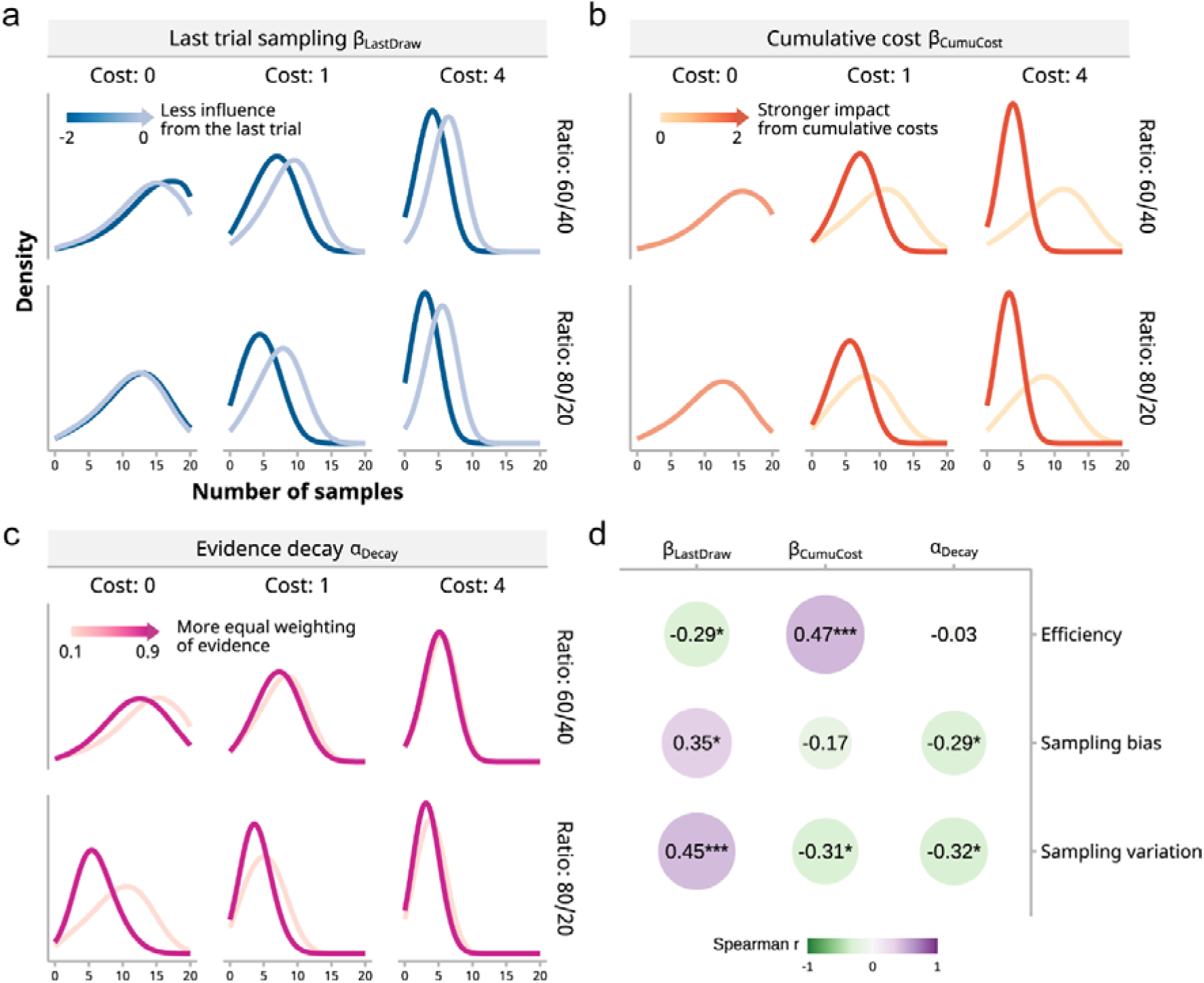
Influence of dynamic decision parameters on sampling behavior. (a) – (b) To illustrate the effect of dynamic parameters, we conducted simulations of children’s sampling processes by holding other parameters constant and varying the interested variables individually within the observed range of the children’s estimates. The influence of β_LastDraw_ and β_CumuCost_ was evident: a stronger impact from the last trial’s sampling and cumulative costs resulted in fewer samples and reduced variation, particularly at higher costs. (c) Simulations for α_Decay_ indicated that a more equal weighting of evidence led to a reduction in the number of samples, particularly under conditions with more evidence and no cost. (d) Correlation between parameter estimates and behavioral metrics: Spearman correlation coefficients reveal the relationships between dynamic parameters (individual-level estimates of β_LastDraw_, β_CumuCost_, α_Decay_) and behavioral metrics (efficiency, sampling bias, and sampling variation). Significance levels are marked by asterisks (\**p* < 0.05, \*\*\**p* < 0.001), with p-values adjusted for false discovery rate (FDR).

Based on the qualitative findings from the simulations, we further conducted correlation analyses between these dynamic parameters and children’s sampling behaviors, and the results exhibited consistent patterns (Figure 5d). β_LastDraw_ showed a significant negative correlation with the average efficiency throughout the experiment (*r*_S_ = -.29, adjusted 95% CI [-.51, -.04], *p* = .028). Such a negative correlation would be even stronger in high-cost conditions (*r*_S_ = -.39, adjusted 95% CI [-.58, -.15], *p* = .002) or in high-evidence conditions (*r*_S_ = -.38, adjusted 95% CI [-.58, -.14], *p* = .006). The direction of these correlations was consistent with the group differences in sampling efficiency and model parameters: autistic children had lower sampling efficiency overall and in high-cost conditions or high-evidence conditions, and their β_LastDraw_ was larger than that of NT children. β_CumuCost_ was positively correlated with the efficiency (*r*_S_ = .47, adjusted 95% CI [.24, .64], *p* < .001), which mirrored the group difference in both the parameter and overall efficiency. Although α_Decay_ was not correlated with the overall efficiency, but as the simulation predicts, it was negatively correlated with the efficiency under the zero-cost conditions (*r*_S_ = -.35, adjusted 95% CI [-.55, -.11], *p* = .021; see Figure 5c left panels).

The direction of the correlations between these parameters and the sampling bias and the sampling variation was also consistent with the group differences. β_LastDraw_ showed a significant positive correlation with the average sampling bias (*r*_S_ = .35, adjusted 95% CI [.011, .56], *p* = .012); and a positive correlation with the average sampling variation (*r*_S_ = .45, adjusted 95% CI [.22, .63], *p* < .001). β_CumuCost_ only had a significant correlation with the sampling variation (*r*_S_ = -.31, adjusted 95% CI [-.52, -.06], *p* = .025). α_Decay_ also had significant negative correlations with sampling bias (*r*_S_ = -.29, adjusted 95% CI [-.50, -.04], *p* = .032) and sampling variation (*r*_S_ = -.32, adjusted 95% CI [-.53, -.08], *p* = .013), which was also compatible with the group differences in α_Decay_ (autistic children had lower α_Decay_ than neurotypical children) and the measures of sampling behavior. These correlations between the estimated parameters and the behavioral measures, combined with the simulations, suggested that autistic children were less flexible and adaptive to the dynamic changes in the environment but more sensitive to the static and local nature of conditions. Such cognitive characteristics of autistic children further lead to their higher variation in sampling decisions, and ultimately relatively poorer sampling efficiency.

## Discussion

The current study adapted the classic bead task into a child-friendly version to investigate differences between autistic and neurotypical children in information sampling with explicit costs. Model-free analyses showed that both groups showed suboptimal sampling but for distinct reasons. Neurotypical children tended to undersample overall. In contrast, the autistic group did not exhibit consistent oversampling or undersampling but instead higher trial-to-trial variability in costly conditions, which resulted in lower sampling efficiency compared to their neurotypical peers. Computational modeling further revealed that both autistic and neurotypical children considered cost- and evidence-related factors in their decision to stop sampling. However, the autistic group, compared to the neurotypical children, were less influenced by last trial sample numbers and cumulative costs but favored recent evidence in a trial. Model simulations and correlational analyses confirmed that these factors, particularly the influence of the last trial samples, contributed to the differences in sampling decision efficiency, sampling bias, and variation observed between the two groups.

Experimental conditions in the current study were divided into distinct blocks, where trials within the same block shared the same information gain and cost, though the samples in different trials were independent of each other. This setting should have facilitated children’s learning of the optimal policy, but autistic children did not exhibit a stable sampling strategy within these blocks. Our modeling results suggest this could result from autistic children’s struggles to effectively use historical and dynamic information to learn and adjust their strategies, coupled with their more weight assigned to recent evidence over earlier evidence in a trial. These findings are reminiscent of the Bayesian account of ASD, including the “hypo-prior” hypothesis [47] and the “high sensory precision” hypothesis [48–50]. According to this framework, autistic people give relatively less weight to prior knowledge or top-down predictions in the inference process, or more weight to sensory inputs, leading to an imbalanced reliance on incoming sensory evidence over prior beliefs during perceptual inference [13, 49, 51] and “continuously high learning rates weakening the influence of past perception on current perception” [14, 50]. From this perspective, our findings of a diminished influence of sample numbers from the previous trial can be regarded as weak prior information for decisions in the current trial, and a stronger weight on recent evidence indicates a higher learning rate in terms of integrating information stream. Our findings also extend this unifying Bayesian theoretical framework for ASD, in the sense that the belief updating in previous research was only about specific percept or value, while ours are also about more abstract sampling strategy.

Further extending these insights, our results also resonate with the Weak Central Coherence (WCC) account of ASD. Traditionally focusing on perceptual processing, WCC theory suggests that autistic individuals tend to focus on details and local information, sometimes at the expense of broader and global context [52–54]. For example, compared to neurotypical individuals, autistic people have shown superior performance in visual search and embedded figures tasks [55], as well as an attenuated McGurk effect, indicative of less integration of visual and auditory cues [56]. Prior research on these theories predominantly addressed perceptual inference [57–66], but both the detail-focused cognitive style and the improper weighting between prior beliefs and new information could manifest in various facets. Our findings align with this detail-oriented cognitive style, where autistic children prioritize recently gathered evidence (i.e., local information) over historical data from previous trials and samples (i.e., global information) in their sampling decisions. Our study expands beyond perception in ASD to delves into information sampling as a form of disambiguatory active inference, which involves individuals actively collecting information from the environment for better inferences in the future [14, 67]. Such actions (i.e., active inference) could link sensory processing differences to a broader range of autistic symptoms, suggesting ASD can be understood as a difference in the balance between perception and action [14]. Thus, our study not only provides new supportive evidence for this perspective but also underscores the importance of extending the Bayesian and WCC accounts of ASD beyond perception to include action. This extension is crucial, given that the core symptoms of ASD are tightly related to the interaction with the world.

Our findings may also shed light on the mixed results found in previous studies on information sampling in ASD. Jänsch & Hare found adults with Asperger syndrome sample less evidence compared to neurotypical adults in a classic “bead task” [19], where participants choose from which bead jar a sequence of beads is more likely to be drawn. Similarly, Farmer et al. found that autistic adults sample less information and make faster decisions, rather than adopting a deliberative sampling style [18]. However, in a similar bead task, Brosnan et al. showed that autistic adolescents gather more beads (i.e., more information) compared to their neurotypical peers [20]. Vella et al. found in a similar box task that autistic participants collect more information to make more correct choices [21]. While the inconsistency might stem from the differences in sample characteristics like age and diagnosis, it is noteworthy that these studies provided limited insights beyond just how much information participants sampled. In particular, the absence of explicit sampling costs or internal cost measurements in these studies makes it hard to assess what role the costs played and how participants responded, because whether oversampling or undersampling is observed depends on the sampling cost, as shown in our results of sampling bias (Figure 2d). Moreover, given our findings that autistic participants base their sampling decisions on a more local context, the seemingly inconsistent observations of oversampling or undersampling in previous studies may arise from subtle differences in their experimental designs.

Age differences also play an important role in information sampling. For example, studies have established distinctive profiles for adolescents, including risk-taking [10, 68–70] and information sampling [2, 5]. These aspects might contribute to the discrepancies observed in Jänsch & Hare and Brosnan et al. [19, 20], where even neurotypical adolescents and adults showed different information sampling behaviors, even under a condition with the same bead ratio and no explicit costs. Comparing our findings with Lu et al. [22], who also used a bead task with explicit costs and multiple ratios, we noted similarities and differences in information sampling between children and adults on the autistic spectrum. Both autistic children and adults with more autistic traits had greater sampling variability under costly conditions, leading to the lower efficiency. However, while our modeling results and those of Lu et al. indicate reduced flexibility in adapting to experimental changes in autistic individuals, the underlying cognitive mechanisms differ between children and adults. We found that autistic children had a cognitive process similar to neurotypical children but were more influenced by static and local factors in the experiment. In contrast, Lu et al. showed adults with more autistic traits adhered to a rigid and fixed strategy. Future research should systematically investigate how information sampling behavior in autistic individuals at different developmental stages is regulated by contextual factors and how it develops. For instance, Crawley et al. [71] demonstrated through computational models that individuals at different age stages, whether they had an ASD diagnosis, used different learning strategies in a probabilistic reversal learning task. This suggests that the learning mechanisms may become more complex as cognitive functions develop. Although direct comparisons between our study and Lu et al. could be challenging due to differences in diagnosis and experimental details, both studies collectively provide strong evidence that variations in information sampling under different information gain and cost conditions may emerge early in autistic individuals and even persist across ages.

In our research, we focused on one type of information sampling, which is instrumental information sampling that by obtaining more information, the probability of making correct decisions can be increased and higher expected returns can be obtained [72, 73]. However, it is also true that the obtained information sometimes cannot change the result of decisions but can only reduce the related uncertainty in advance. This type of information sampling is called non-instrumental information sampling [72, 74–76]. Studies have suggested that the intrinsic value of non-instrumental information may partly be related to ameliorating negative affects caused by uncertainty [74, 77, 78], which makes non-instrumental information highly relevant to autism. Research has found that handling uncertainty about events is challenging for individuals with autism and often leads to anxiety [79–81]. Some theories also suggest that such anxiety may induce common restricted and repetitive behaviors in autism, such as shifting attention to limited interests, sensory-seeking, or repetitive behaviors to regulate and attenuate the anxiety [82]. Therefore, it is crucial for future research to investigate how autism affects non-instrumental information sampling behavior and the relationship between non-instrumental information sampling and repetitive behaviors in autistic people.

### Limitations

Though all the autistic children in our study had a formal ASD diagnosis by professional clinicians, we enrolled the participants from different waves and sites, leading to a situation where children, particularly autistic children, underwent additional assessments that served similar functions but were different in the form (e.g., among the 24 autistic children, 15 children had ADOS, 5 children had CARS and AQ-child measures, and the remaining 4 children had AQ-child measures). In addition to the formal ASD diagnosis, those assessed with ADOS or CARS met the respective cutoffs, and those tested with AQ-child exhibited significantly higher scores than the neurotypical children (Table 1). Nevertheless, this inconsistency in assessment tools prevents us from adopting a dimensional approach to correlate our modeling results with autistic symptom severity, as recently advocated in psychiatry [83, 84]. A second limitation is the sample size (i.e., the number of participants). The sample size of our study lines up with those of recent related studies on decision-making, learning, and inference in autistic children or adults [61, 65, 85–93], leading to robust findings that accorded with those found in adults with varying autistic traits [22]. However, if we had had resources to test more participants (the high functional autistic children is relatively rare), we would be able to refine subtyping based on our computational model parameters, which could offer more insights into each individual and ASD per se [94–96]. Future studies could benefit from a larger sample size and more comprehensive assessments to achieve not only greater statistical power but also deeper insights into individual and subgroup differences, hopefully contributing more to precise interventions.

## Conclusion

In summary, we show that autistic children perform differently in an information sampling task compared to neurotypical children of the same age and intelligence level. Using computational modeling, we find that autistic children are less affected by dynamic changes between and within trials and more inclined to weigh recent pieces of evidence, resulting in greater behavioral instability and lower sampling efficiency. The information sampling behaviors of autistic children resemble those of adults with more autistic traits, despite the strategic differences probably due to ages. These findings reveal developmental differences in a fundamental cognitive function at an early stage, and may provide empirical support for and even expand the Bayesian framework of ASD.

## Declarations

### Ethics approval and consent to participate

The Institutional Review Board of the School of Psychological and Cognitive Sciences at Peking University approved the study, and all the child participants and their parents provided informed consent according to the Declaration of Helsinki.

### Consent for publication

Not applicable

### Availability of data and materials

The codes and datasets generated and analyzed during the current study are available from the corresponding author on reasonable request and in accordance with ethical procedures for the reuse of sensitive data.

## Authors’ contributions

HL collected, analyzed, and interpreted the data for statistical analyses and computational modeling, and was a major contributor in drafting the manuscript. HZ and LY provided supervision and funding acquisition. All authors participated in conceptualization and reviewing and editing the manuscript.

## Acknowledgements

We would like to thank all the children and their parents for participating in our study. We are also thankful to Tianbi Li, Yixiao Hu, Zheng Wang, and staff in Qingdao Elim School for their generous assistance for the study.

