## Additional File 1 for "Beyond over- or under-sampling: autistic children’s inflexibility in sampling costly information"

### S1 Text: Supplementary methods

#### Task instruction and comprehension check questions

Note that the task instruction and the comprehension questions were asked in the language from the experiment site.

**Task instructions.** "Today, you're a little explorer who will go on an adventure to either Doggy Island or Kitty Island. Look, there are more doggies on Doggy Island and more kitties on Kitty Island. Your task is to figure out whether you're on Doggy Island or Kitty Island. On the island, you'll meet some small animals. Look here, you've encountered two cats and one dog. These three animals might come from Doggy Island or Kitty Island. If you think there are more kitties than doggies now, where do you think we are more likely to be – on Kitty Island or Doggy Island? Since there are more kitties on Kitty Island, it's more likely that we'll meet more kitties there. Besides adventuring in winter, we'll also go on adventures in summer. Look, during summer, the difference in the number of doggies and kitties is even smaller, making it harder to guess. Look at the top of this screen, there are 100 cookies to feed the kitties and doggies we meet. We might meet three types of animals. First, are the tiny animals. They are so small that we don't need to feed them cookies. No matter how many tiny animals we meet, the number of cookies won't decrease. We may also meet animals of regular size. For each regular-sized animal we meet, they will eat one cookie, like now, we've met one and there's one less cookie. The more of these animals we meet, the fewer cookies we'll have left. Finally, we might meet big animals. Big animals are so large that they eat four cookies each time we meet one. So, the more big animals we meet, the quicker our cookies will disappear. You will only get coins as a reward if you successfully determine whether you're on Doggy Island or Kitty Island. Sometimes, you might not guess correctly, and then you won't get any coins. But that's okay because unexpected things can happen. Just try to make as many successful guesses as you can. The more coins you get, the more sticker rewards you'll receive. So, in the game, you need to think and decide when you're confident enough to guess where you are based on how many animals you meet, and always pay attention to how your cookies change."

**Comprehension check questions.**

1. “When you have encountered these three animals, at which island do you think you are more likely to meet them?” (Children were asked for each of the following situations, thus four questions in total.)

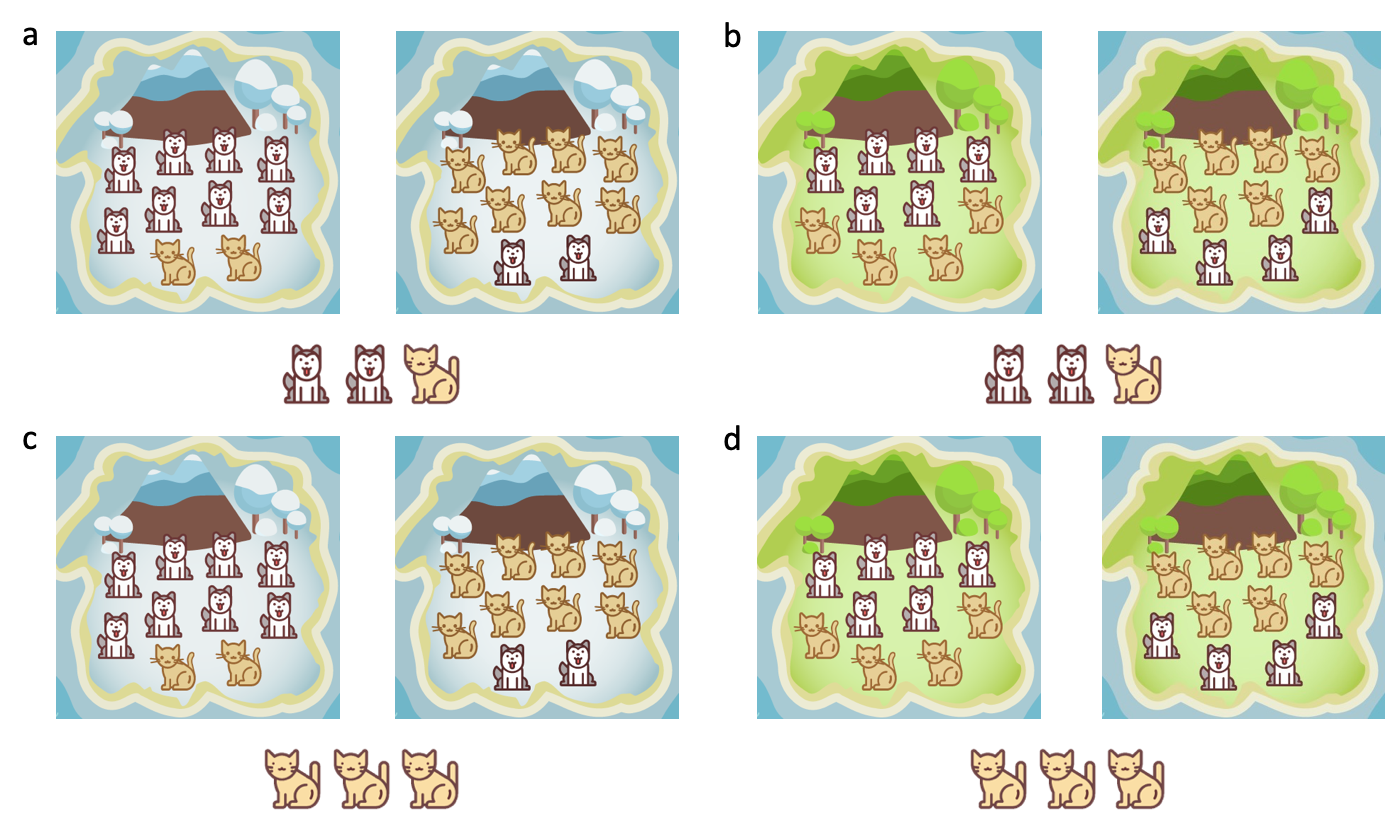

2. “How many cookies do one tiny/regular/big animal eats? If you meet many tiny/regular/big animals, what will happen to your cookies? Will they decrease?” (Children were asked about each type of animals, thus three questions in total.)

#### Parameter specifications of hierarchical Bayesian models

We assume that a generic individual-level parameter, say β, was drawn from a group-level normal distribution; that is, β ~ Normal(*μ*_β_, *σ*_β_), with *μ*_β_ and *σ*_β_ being the group-level mean and standard deviation, respectively. Both these group-level parameters were specified with generic weakly informative priors, following the recommendation of Stan Dev team [1]: *μ*_β_ ~ Normal(0, 1) and *σ*_β_ ~ Half-t(4, 0, 0.5). Since the number of groups was small specifically in our autistic sample, the data might have a chance not providing much information on the group-level variance, so we chose a relatively stronger prior information on the scale parameter (i.e., the standard deviation) that allowed more pooling. Several parameters were constrained within [0, 1] (with inverse logit transform), including decay parameter α_Decay_ in models with decayed evidence and second-thought probabilities.

### S1 Table: Model descriptions

| Model | One- or two-stage? | Cost-first or evidence-first? | Second-thought probability | Cost-related decision variables | Evidence-related decision variables | N individual-level parameters |
| --- | --- | --- | --- | --- | --- | --- |
| Cost only | One | N/A | N/A | A constant, unit cost (three levels with treatment coding), the number of beads sampled, total sampling cost | NA | 5 |
| Cost + Evidence | One | N/A | N/A | As above | Unit log evidence, absolute value of **decayed** cumulative information, total **decayed** log evidence, last trial sample numbers, last trial correctness | 11 |
| 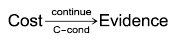 | Two | Cost-first | Controlled by cost conditions (three parameters) | As above | As above, plus a constant | 15 |
| 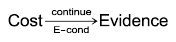 | Two | Cost-first | By evidence conditions (two parameters) | As above | As above | 14 |
| 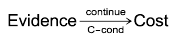 | Two | Evidence-first | By cost conditions | As above | As above | 15 |
| 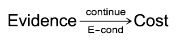 | Two | Evidence-first | By evidence conditions | As above | As above | 14 |

### S2 Table: Summary statistics of group-level mean parameters of the best-fitting model

|  |  | **Group-level mean** | | |
| --- | --- | --- | --- | --- |
|  | Parameter | Autistic group^1^ | Neurotypical group^1^ | Difference^1^ |
| (Constant) | (Constant) | -3.69 [-4.35, -2.98] | -2.95 [-3.48, -2.37] | -0.74 [-1.63, 0.13] |
| Cost-related | Sample numbers | 0.91 [0.35, 1.47] | 1.66 [1.03, 2.26] | -0.75 [-1.59, 0.07] |
|  | Low- to zero-cost | 1.07 [0.54, 1.61] | 0.3 [-0.03, 0.64] | 0.77 [0.16, 1.41] |
|  | High- to zero-cost | 0.77 [0.21, 1.32] | -0.06 [-0.54, 0.44] | 0.83 [0.11, 1.59] |
|  | Cumulative cost | 0.72 [0.47, 0.97] | 1.2 [0.79, 1.59] | -0.48 [-0.95, -0.01] |
| Evidence-related | Unit log evidence | 0.18 [-0.03, 0.38] | 0.27 [0.15, 0.39] | -0.09 [-0.33, 0.14] |
|  | Last trial correctness | 0.07 [-0.01, 0.14] | 0.14 [0.07, 0.21] | -0.07 [-0.18, 0.03] |
|  | Last trial sample numbers | -0.42 [-0.53, -0.3] | -0.96 [-1.18, -0.73] | 0.54 [0.28, 0.78] |
|  | Absolute cumulative information | 0.03 [-0.62, 0.73] | 0.71 [0.46, 0.97] | -0.68 [-1.41, 0.03] |
|  | Cumulative log evidence | 0.06 [-0.15, 0.27] | -0.07 [-0.14, -0.01] | 0.13 [-0.08, 0.35] |
|  | Evidence decay^2^ | 0.26 [0.05, 0.63] | 0.76 [0.47, 0.94] | -0.46 [-0.81, -0.04] |
| ^1^Median [95% HDI]; ^2^Inverse logit transformed to [0, 1] | | | | |
